# Direct identification of *de novo* mobile element insertions from single molecule sequencing of human sperm

**DOI:** 10.1101/2025.10.25.684559

**Authors:** Stacy Li, Landen Gozashti, Caitlin Connelly, Clément Goubert, Kenneth Aston, Joseph G Gleeson, Aaron Quinlan, Xiaoxu Yang, Peter H Sudmant

## Abstract

Mobile element insertions (MEIs) are a significant source of human genetic variation, yet the rates and properties of *de novo* MEIs are poorly characterized due to technical limitations in sequencing technology. Here, we directly sequenced individual gametes from sperm samples of 19 donors (aged 27-62) using highly accurate PacBio long-read sequencing to identify *de novo* retrotransposition events without familial inference. We developed a “self-alignment” strategy using personalized genome assemblies that enables high-precision, single-read detection of *de novo* MEIs. Using this method, we identified 43 *de novo* Alu insertions, revealing >9-fold variation in Alu retrotransposition rates between individuals (ranging from 0 to 0.148 insertions/gamete). We found a significant increase in Alu activity with paternal age, yielding a 4.67% increase in insertions per gamete per year of additional paternal age, representing a direct observation of age-associated increases in structural variant (SV) mutation rates. *De novo* Alu insertions predominantly represent evolutionarily young AluYa5 and AluYb8 subfamilies and bear characteristic molecular signatures of target-primed reverse transcription (TPRT). Our population-averaged rate of 4.52 insertions per 100 gametes aligns well with previous population genetic estimates, validating both direct observation and population approaches for estimating *de novo* MEI rates. These results establish direct gamete sequencing as a powerful method for characterizing germline mutation processes and reveal age as a significant determinant of *de novo* retrotransposition in the male germline.

## Introduction

*De novo* mutations (DNMs) originate from inaccuracies that occur during parental gametogenesis. These mutations, arising from errors in DNA replication, repair, and recombination, range in size from single base pair substitutions to megabase-scale rearrangements and aneuploidies. DNMs are not uniformly distributed throughout the genome: clusters of DNMs occur more frequently at recombination hotspots, and recurrent genic DNMs underlie multiple neurodevelopmental disorders (Wilfert et al. 2017; Francioli et al. 2015; Hinch et al. 2023). Additionally, the number of genome-wide single nucleotide variant (SNV) DNMs steadily increases with parental age. Phasing of DNMs reveals that the majority of SNV DNMs originate from the paternal gamete. Together, this paternal age effect yields an increase of 2 additional *de novo* single nucleotide variants (SNVs) per year of paternal age at conception.

Even correcting for the increased number of meioses incurred through the male reproductive lifespan, the male germline mutation rate still increases with age (Kong et al. 2012; Jónsson et al. 2017; Goldmann et al. 2019). Comparative analysis of DNMs across amniotes attributes the paternal mutation bias to differences in endogenous DNA damage and DNA repair response, which are further magnified in aging (de Manuel et al. 2022; Kong et al. 2012).

Estimates of DNM rates have almost exclusively been derived from short-read whole-genome sequencing of small variants, leaving a gap in our understanding of *de novo* structural variants (SVs). Short-read analysis of more than 2000 trios with autism spectrum disorder (ASD) found a significant increase in dnSVs in probands compared to unaffected siblings attributed to mutations borne by the paternal gamete (Belyeu et al. 2021). However, short reads often fail to capture larger *structural variants* (SVs) and variants occurring in highly identical or repetitive regions of the genome (e.g. centromeres). As a result, the true rate of *de novo* SVs (dnSVs) is systematically underestimated despite their relevance in human disease.

Long-read sequencing improves both recall and sensitivity of SV discovery, vastly expanding the landscape of genetic variation that can be assayed. For example, DNM discovery in family studies is significantly improved by long-read data, increasing the *de novo* SV yield by over 30% (Noyes et al. 2022). More recently, analyses of T2T genomes identified *de novo* mutations in several previously inaccessible regions of the genome including STRs, centromeres and the Y chromosome (Porubsky et al. 2025). However, family studies can only assess inherited mutations which are compatible with life. Furthermore, family studies are inherently limited by the number of offspring assessed, with each individual representing the product of only two independent parental meioses.

Recent work has shown that some *de novo* SVs, such as transposable element insertions, can be detected in single molecule long-reads, presenting an opportunity to study *de novo* SVs without confounding factors and challenges of family studies (Gerdes et al. 2022; Movilli et al. 2025). Transposable elements (TEs) are repetitive sequences capable of moving throughout the genome. NonLTR retrotransposons, a subclass of TEs, comprise over 25% of the human genome (Hoyt et al. 2022). Amplification and mutation of nonLTR retrotransposons underlie both local-scale instability and long-range evolutionary innovations in genome sequence and structure (Bailey et al. 2003). Ongoing TE mobilization in humans is limited to three families of nonLTR retrotransposons: Alu, LINE-1 (L1), and SVA elements (Cordaux and Batzer 2009). For all of these, listed in order of activity, retrotransposition depends on the activity of ORF2p, a combined endonuclease (EN) and reverse transcriptase encoded by autonomous L1 retrotransposons (Feng et al. 1996; Dewannieux et al. 2003). ORF2p operates as part of a ribonucleoprotein complex, binding ORF1p (a RNA chaperone, co-translated with ORF2p) and a single compatible transcript. ORF2p nicks genomic DNA at a 5’-TTTT/AA-3’ motif (Jurka 1997; Thawani et al. 2024). Cleavage by ORF2p exposes a single-stranded 3’ poly(T) sequence used to prime reverse transcription from the poly(A) tail of the retrotransposon transcript, followed by a second strand nick to prime second-strand synthesis and integration of the template retrotransposon into the genome. This process, called target-primed reverse transcription (TPRT), results in target site duplications (TSDs) flanking the new retrotransposon insertion (Beck et al. 2010; Batzer and Deininger 2002; Luan et al. 1993; Richardson et al. 2017).

Human nonLTR retrotransposons can be further divided into subfamilies on the basis of sequence identity and phylogenetic age estimates. Most ancient sequences have diverged and become immobile, whereas younger copies still capable of mobilizing are frequently polymorphic and linked to increased risks of disease and genomic instability (Payer et al. 2017; Nazaryan-Petersen et al. 2016). Only a fraction of young elements, known as *source elements*, remain mobile in the human genome (Cordaux et al. 2004; Hoyt et al. 2022; Mills et al. 2007). Theoretical and pedigree-based studies suggest that the majority of *de novo* Alue retrotransposition events occur in the male germline (Feusier et al. 2019; Jurka et al. 2002).

However, studies of other non-LTR retrotransposons have identified a female germline bias. Furthermore, mobilization of source elements in the germline is associated with both rare and recurrent genetic disease (Hancks and Kazazian 2012). Together, this class of genetic variation is important for both human disease and evolution.

Here, we sought to directly estimate the properties and rates of *de novo* Alu element insertions in the male germline by using highly accurate long-read sequencing to directly sequence gametes. Alu insertions are a medically relevant class of mutation with a greater potential for deleterious effects compared to single nucleotide polymorphisms (Stewart et al. 2011). New Alu insertions are estimated to occur in approximately 1 out of every 20 to 100 live births (Konkel et al. 2015; Feusier et al. 2019; Werling et al. 2018; Cordaux et al. 2006a; Xing et al. 2009; Deininger and Batzer 1999). As a result of their relatively low frequency, *de novo* Alu activity is challenging to study through conventional pedigree-based approaches; for example, recent analyses of a four generation pedigree identified just a single *de novo* SVA element insertion and no Alu insertions (Porubsky et al. 2025).

Here we develop a *self-alignment* based (i.e. personalized genome-based) strategy that improves variant calling precision by reducing reference bias through the use of sample-specific genome assemblies. This approach enables reliable detection of *de novo* events supported by single reads, facilitating direct observation of active mutational processes in the germline.

## Results

### A framework to identify de novo retrotransposition events in single reads

Conventional germline variant callers rely on support from multiple reads to confidently detect polymorphisms (Poplin et al. 2018; Heller and Vingron 2019). However, these approaches are less suitable for non-germline variants, such as somatic and *de novo* variants. These variants occur at low allelic fractions (AF), making them challenging to distinguish from sequencing error, especially in samples where allelic heterogeneity is expected (e.g. tumors). Numerous strategies are employed to optimize sensitivity and specificity in detection of low-AF variants. One common approach reduces erroneous calls by comparing candidate variants against a reference set of variants (e.g. matched tumor-normal variant calling). Another common approach, ensemble variant calling, identifies high-confidence variants by intersecting multiple callsets. Inspired by these techniques, we developed a novel approach for directly identifying and phasing *de novo* variants in gametes using long reads without a parental reference.

We performed PacBio HiFi long-read sequencing on bulk sperm samples from 19 individual donors, including four samples from donors with children affected by TSC (tuberous sclerosis complex) or ASD (autism spectrum disorder) **(Fig. 1A-B)**. We sequenced each sample to an average depth of ∼50.9 (Supplementary Table 1; see Methods), capturing between 1 million and 2.5 million unique cells per individual. We estimate that the vast majority of reads are unique products of a single meiosis and each cell is covered by at most 4 reads (see **Methods**).

**Figure 1:**
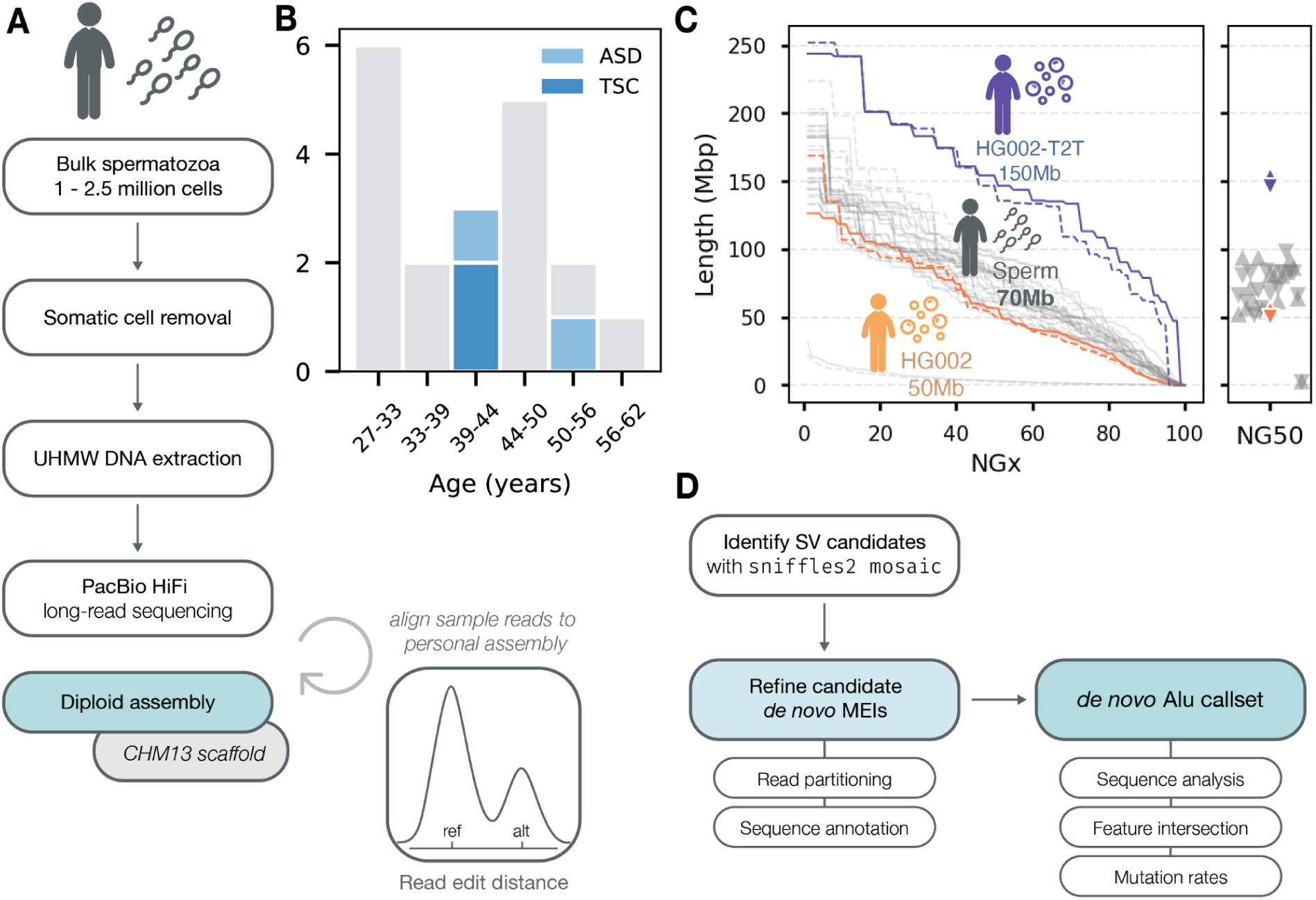
Identifying *de novo* retrotransposition events from individual sperm cells. Experimental design and assembly quality metrics. **A)** Workflow for DNA extraction, sequencing, genome assembly, and self-alignment. Bulk spermatozoa samples ranging from 1-2.5 million cells were obtained from 19 donors. Ultra-high-molecular-weight (UHMW) DNA was extracted after somatic cell removal. We performed PacBio High Fidelity (HiFi) sequencing and generated a diploid personalized assembly for each sample, which we then scaffolded to CHM13. Aligning raw reads from each sample to its personalized reference minimizes edit distance between non-SV bearing reads and the reference. **B)** Histogram of donor age. **C)** Assembly contiguity comparison between 19 sperm-derived assemblies and reference datasets, showing comparable N50 values. **D)** Variant calling pipeline using self-assembly alignment and ensemble methods. We identified SV candidates using sniffles2 mosaic and refined *de novo* mobile element insertions (MEIs) through extensive filtering (see Methods).

Next, we generated phased diploid assemblies for each individual using *hifiasm (Cheng et al. 2021)*, yielding an average NG50 of ∼71 Mb per haplotype. The majority of sperm-derived haplotype-resolved assemblies were comparable in contiguity and quality to an assembly generated from a diploid control dataset (HG002 HiFi reads from HPRC, B-lymphocyte) (Liao et al. 2023) (**Fig. 1C**). We then aligned reads from each sample to its matched genome assembly (“self-alignment”). This “self-alignment” approach increases precision by minimizing the edit distance between non-SV bearing reads and the reference genome, enabling us to distinguish true *de novo* events from alignment artifacts and allelic variation present in the donor individual.

Following alignment, we performed variant calling in two phases **(Fig. 1D)** (see Methods for full details). Briefly, variant candidates were called using sniffles2 (Smolka et al. 2024) with the low-frequency “mosaic” variant detection mode, optimizing for sensitive detection of variants occurring in single reads by removing default quality control filters. We then filtered this first set of variants to only include low-frequency insertions, all of which except one occurred within a single read. In the second phase, insertion sequences were annotated using GraffiTE’s repeat filtering and target site duplication modules to identify high-quality candidate retrotransposition events (Groza et al. 2024). These candidates were then subjected to stringent filtering for hallmark features associated with *de novo* TE insertions such as identical flanking target site duplications, optimizing for precision (see **Methods**). Finally, we filtered potential false positives caused by mapping errors and contamination by remapping candidate *de novo* insertions and their flanking context back to the self assembly and reference genome. We also checked for intersample contamination which might yield false positives (see **Methods**).

### Self-assembly alignment improves variant calling precision

To benchmark our approach we developed *nova*, a simulation framework that introduces predefined variants into individual reads from a reference dataset. We generated thousands of random synthetic *de novo* MEIs for both Alus and L1s as well as a random set of non-MEI-related insertions of various sizes from reads sampled equally across our 19 sperm donors. We used the synthetic de novo MEIs as positive controls and other insertions as negative controls for pipeline evaluation. Then, we performed alignment with two strategies: (1) a standard approach with alignment to the CHM13 reference genome, and (2) a self-alignment approach to each diploid self-assembly (Jarvis et al. 2022). Variant calling against CHM13 yielded 0 true positives for both Alus and L1s, whereas our self-alignment exhibited a false negative rate of only 0.03 and 0.02 for Alus and L1s respectively (Supplementary Figure 1).

These results suggest that self-alignment is necessary for sensitive *de novo* MEI detection. We also tested the performance of sniffles2 mosaic against another recently developed mosaic variant caller minisv, which integrates both self-assembly as well as a reference genome to improve variant calling (Qin et al. 2025). Sniffles2 exhibited a sensitivity of 0.96 and accuracy of 0.98 across all simulated de novo MEIs (**Fig. 2A-B)**. While miniSV demonstrated comparable accuracy (>0.94), it was far less sensitive compared to sniffles2 (**Fig. 2A-B**). This was especially the case L1s, where sensitivity was only 0.62 (**Fig. 2B)**. Thus, we implemented sniffles2 in our finalized workflow. Notably, our pipeline produced zero false positives in all benchmarks, irrespective of the upstream variant caller or reference, suggesting robust automated curation of prospective *de novo* MEIs. Together, these results demonstrate that *de novo* MEIs can be accurately called from single reads mapped to a self-assembly.

**Figure 2:**
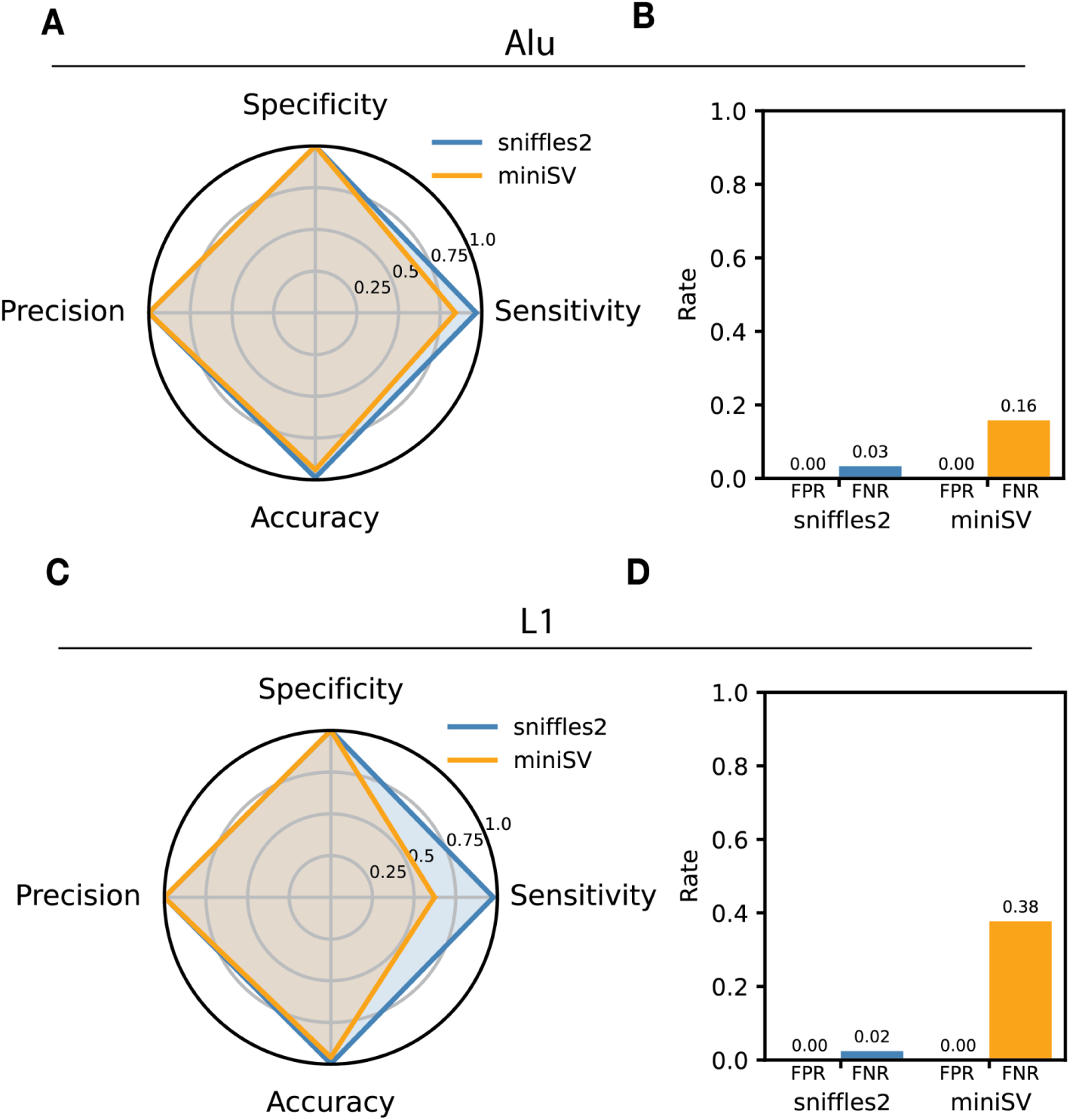
Simulation and detection of synthetic single read mobile element insertions. *nova* simulates *de novo* MEIs by sampling and modifying reads from a dataset of long reads. **A**) Radar plot comparing precision, specificity, accuracy, and sensitivity across sniffles2 and miniSV approaches for simulated *de novo* Alu insertions. **B)** False positive rates (FPR) and false negative rates (FNR) for simulated *de novo* Alu insertions. **C)** Radar plots comparing sniffles2 and miniSV variant calling approaches for simulated *de novo* L1 insertions. **D)** FPR and FNR for simulated *de novo* L1 insertions.

### Direct identification of *de novo* mobile element insertions from long reads

We next applied our approach to identify *de novo* Alu retrotransposition events from individual sperm samples. We identified a total of 43 high-confidence full-length *de novo* Alu insertions in single long reads, capturing both the Alu sequence and the genomic context of the insertions (Supplementary Table 2). We identified characteristic features of each Alu, including the insertion orientation, poly(A) tail, and TSD for each insertion **(Fig. 3B)**. We find nearly equal numbers of sense and antisense oriented Alu insertions (53.5% sense and 46.5% antisense), reflecting genome-wide proportions of Alu orientation (Cook et al. 2011). We also identified a single, truncated L1 insertion on chr4 (Supplementary Table 2). For the remainder of the manuscript however we focus on the properties of the Alu insertions.

**Figure 3:**
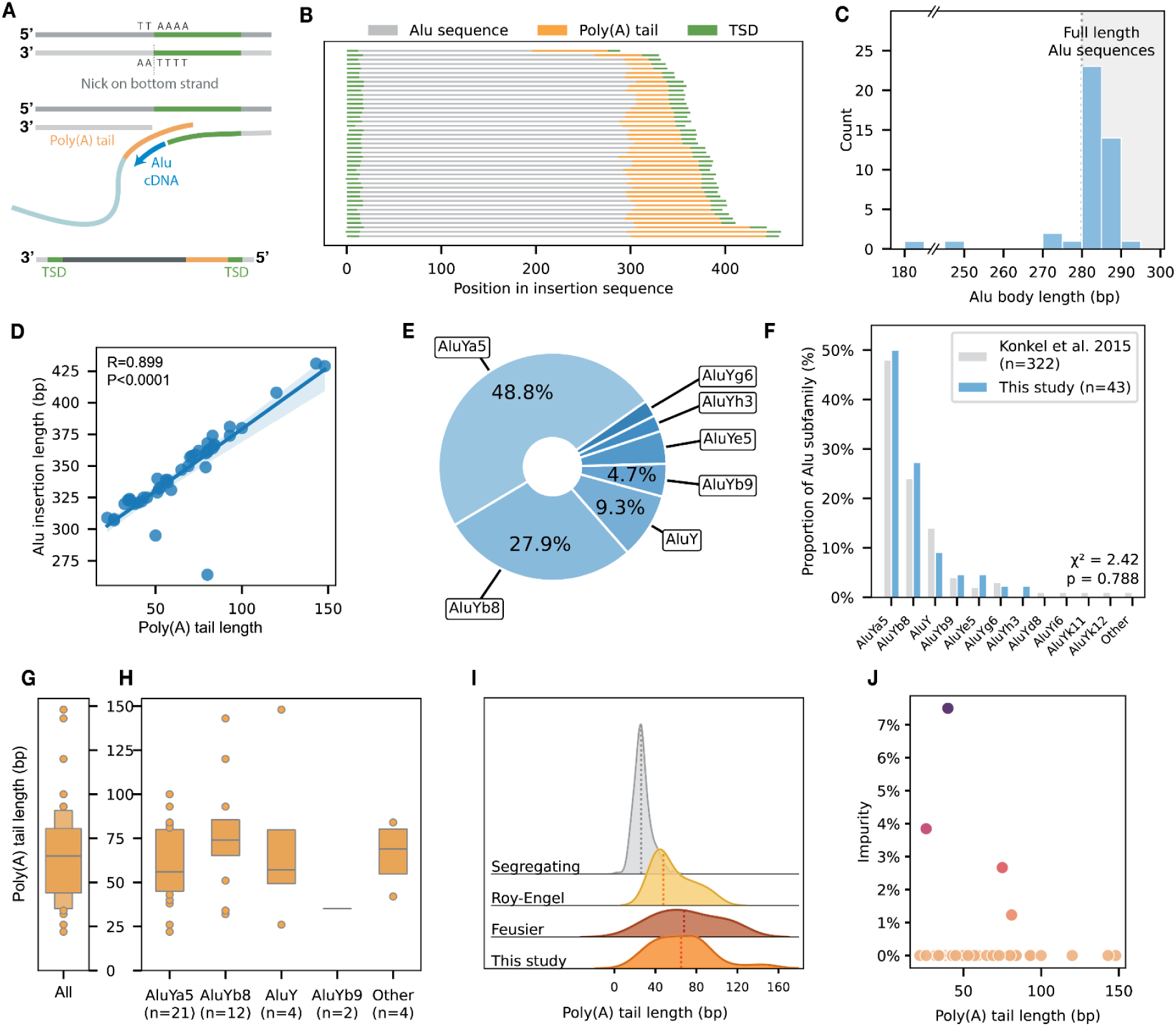
Molecular features of de novo Alu insertions. **A)** An illustration of target-primed reverse transcription ofAlu elements. ORF2p cleaves the top or bottom (bottom shown) strand at a 5’-TTTT/AA-3’ L1 EN recognition motif. Cleavage of the “bottom” strand yields a *sense* Alu and duplication of the target site region on the 5’ end of the bottom strand. **B)** Graphical representation of 43 discovered *de novo* Alu insertions and their characteristic features. Flanking target site duplications in green. **C)** Histogram of Alu body lengths (Alu sequence minus the poly(A) tail). The interval highlighted in gray indicates the range of full length Alu body sequences. **D)** Correlation between total Alu insertion length and poly(A) tail length with 95% confidence intervals (Spearman correlation, R = 0.899, P<0.0001). **E)** Subfamily composition of *de novo* Alus. All identified Alus are members of the AluY subfamily and its derivatives. AluYa5 and AluYb8 are disproportionately mutagenic compared to other subfamilies. **F)** Comparison of *de novo* Alus and segregating Alus analyzed in the 1000 Genomes Project. Segregating Alus originate from *de novo* Alu mobilization events in the recent evolutionary past. The distribution of subfamilies represented in the 1000 Genomes data closely matches the distribution of *de novo* insertions in our study. **G, H)** Boxenplot distribution of poly(A) tail lengths in total and across subfamilies for *de novo* Alu insertions. The median length (56 nt) is marked by the center line dividing the upper and lower quantiles. Outliers are shown in points above and below the range of the quantile boxes. **I)** Kernel density estimation plot of *de novo* Alu poly(A) tail lengths in this study compared to *de novo* Alu insertions in two previous studies (Roy-Engel et. al 2002: n = 16; Feusier et al. 2019: n = 11) as well as 37 young segregating Alus (Roy-Engel et al. 2002). The dotted lines represent the median poly(A) tail lengths for each category (from top to bottom: 26 nt, 48 nt, 68 nt, 65 nt). **I)** Pol(A) tail impurity, defined as the percent of non-A nucleotides in a sense poly(A) tail.

Most Alu insertions are full length, spanning the main body of the Alu sequence (280bp) and terminating in a poly(A) tail of variable length, noting that we required a minimum insertion length of 250bp during variant filtering (**Fig. 3C**). Nonetheless, total lengths (body length plus the poly(A) tail) varied considerably across Alu insertions (min = 264, max = 431, median = 340; **Fig. 3B**, Supplementary Figure 2). We found that poly(A) tail length variation explained >80% of the variance in *de novo* Alu insertion lengths, with the longest insertion exhibiting a remarkable 148bp poly(A) tail (**Fig. 3D**; Spearman correlation, R=0.899, P<0.00001). We also observed a small minority (∼9%, 4/43) of sequences with 5’ truncations ranging from 8 to 96 bp **(Fig. 3B,C)**. This proportion is similar to the proportion of 5’ truncated Alus identified from population-scale sequencing of segregating Alus in the 1000 Genomes Project (two-sided Binomial test, P = 0.510) (Konkel et al. 2015).

### Segregating Alu subfamilies are representative of *de novo* Alu subfamilies

Mutations accumulated in Alu source elements are inherited by descendant copies, resulting in Alu lineages with variable sequence content and activity. Phylogenetic analysis of these elements partitions human-specific Alus into three major groups: the evolutionarily ancient AluJ and AluS subfamilies, and the more recent AluY subfamily (Liu et al. 2009; Batzer and Deininger 2002). Analysis of segregating Alus suggests that only AluY-derived elements remain mobile in the human genome (Konkel et al. 2015). Consistent with this, we find that all 43 of our *de novo* Alu elements belong to AluY and its derivative subfamilies, with AluYa5 and AluYb8 together comprising over 75% of identified insertions (**Fig. 3E**). The frequency of *de novo* insertions across AluY subfamilies in the germline is also consistent with the population genetically identified Alu profile established by the 1000 Genomes Project (p = 0.788: chi-square goodness of fit) (**Fig. 3F**) (Konkel et al. 2015).

### Sequence features of *de novo* Alu insertions

### De novo Alu insertions are marked by significantly elongated poly(A) tails

Experimental dissection of Alu activity has identified a direct association between the length of the poly(A) tract of the RNA and transposition frequency (Dewannieux and Heidmann 2005a; Comeaux et al. 2009). Alu sequences require a poly(A) tail of at least 10-15 nt to mediate cDNA synthesis during TPRT to mobilize, with each additional 10 nt contributing to a logarithmic increase in activity, plateauing beyond 50 nt (Comeaux et al. 2009; Dewannieux and Heidmann 2005b). Furthermore, following integration into the genome, progressive degradation of poly(A) length and sequence content occurs leading to eventual inactivation of Alu elements (Dewannieux and Heidmann 2005a; Roy-Engel et al. 2002). This is supported by the observation of relatively shorter A-tails in ancient, immobilized Alus (average: 21 nt) compared to those found in relatively new, segregating Alus (average: 29 nt) (Konkel et al. 2015; Roy-Engel et al. 2002).

We leveraged the extended context of each supporting read to characterize the poly(A) tail length and content of each Alu, with the caveat that homopolymers can have a higher error rate in PacBio reads (Ono et al. 2022). We identify a wide distribution of poly(A) tail lengths strongly skewed towards extended poly(A) tracts (min: 22 nt, median: 65 nt, max: 148 nt) (**Fig. 3G-H).** We find a significant difference between segregating and *de novo* A-tail lengths, skewed longer in *de novo* insertions (p < 0.001; Mann-Whitney U test). Two previous studies using Sanger sequencing and short-read sequencing identified similar A-tail length distributions in pathogenic *de novo* Alus, as well as a similarly significant difference between segregating and *de novo* A-tails (**Fig. 3I**) (Dewannieux and Heidmann 2005a; Roy-Engel et al. 2002; Feusier et al. 2019). We observe a greater maximum tail length but no significant difference in median lengths between our study and previous studies of *de novo* events (p = 0.405; Kruskal-Wallis test), highlighting the improved sequence capture afforded by long-read sequencing.

Next, we examined the purity of each poly(A) sequence. The majority of A-tails were entirely homopolymeric, consistent with *de novo* insertions generated by mobile source elements. We observed several notable exceptions bearing unusually high frequencies of non-A bases in below-median tail lengths (**Fig. 3J**; Supplementary Table 2). Manual examination revealed single-base impurities interspersed within the poly(A) sequence, rather than contiguous tracts of non-A bases. Notably, within individual sequences, all impurities consisted of the same nucleotide (e.g., all T or all G). We hypothesize that these impurities are likely the result of PacBio sequencing errors in homopolymer tracts (Ono et al. 2022), although rapid accumulation of poly(A) tail impurities has been reported in somatic tissues (Evrony et al. 2015). Together, our analysis supports a model in which poly(A) tail length increases upon transcription of a mobile element, then shortens and degenerates progressively after insertion.

### Target sites and target site duplications

We analyzed the sequence context at insertion breakpoints to identify motifs associated with Alu insertions. In the immediate context surrounding the breakpoints, we identified the canonical 5’-TTTT/AA-3’ ORF2p endonuclease recognition motif (Supplementary Figure 3) (Flasch et al. 2019; Szak et al. 2002; Konkel et al. 2015). ORF2p-mediated TPRT yields TSDs between 6 and 22 nt that are enriched in adenosine content at the proximal positions of the insertion motif region (Szak et al. 2002; Konkel et al. 2015; Ewing et al. 2020). We identified TSDs between 9-18 nt (median: 15 nt) in length in all 43 Alu insertions (Supplementary Figure 4). TSD lengths of *de novo* Alu insertions are less variable compared to segregating Alus reported in the 1000 genomes project (Stewart et al. 2011) (Levene’s test; P=0.0053). However we note that this could be due to methodological differences in how they are ascertained. Moreover, the presence of TSDs is a requirement for *de novo* MEI discovery in our pipeline.

### Sequence preferences of Alu insertions

Next, we sought to characterize the genomic context and distribution of *de novo* Alu insertions. *De novo* Alu insertions are distributed across most human chromosomes (18/24, **Fig. 4A**). Chr19 exhibits the highest density of segregating and fixed Alus among human chromosomes as well as the highest gene density (Grover et al. 2004a) have been largely shaped by selection, but could also result from insufficient sampling. Furthermore, previous studies have suggested that Alus may preferentially nest within other Alus or L1 elements (Levy et al. 2010). To test for this, we annotated mobile elements and other repeats in all self-assemblies using RepeatMasker. We compared the proportion of de novo Alu insertions in different repeat types to expectations from randomly resampled degenerate L1 endonuclease motif coordinates (Flasch et al. 2019).

**Figure 4:**
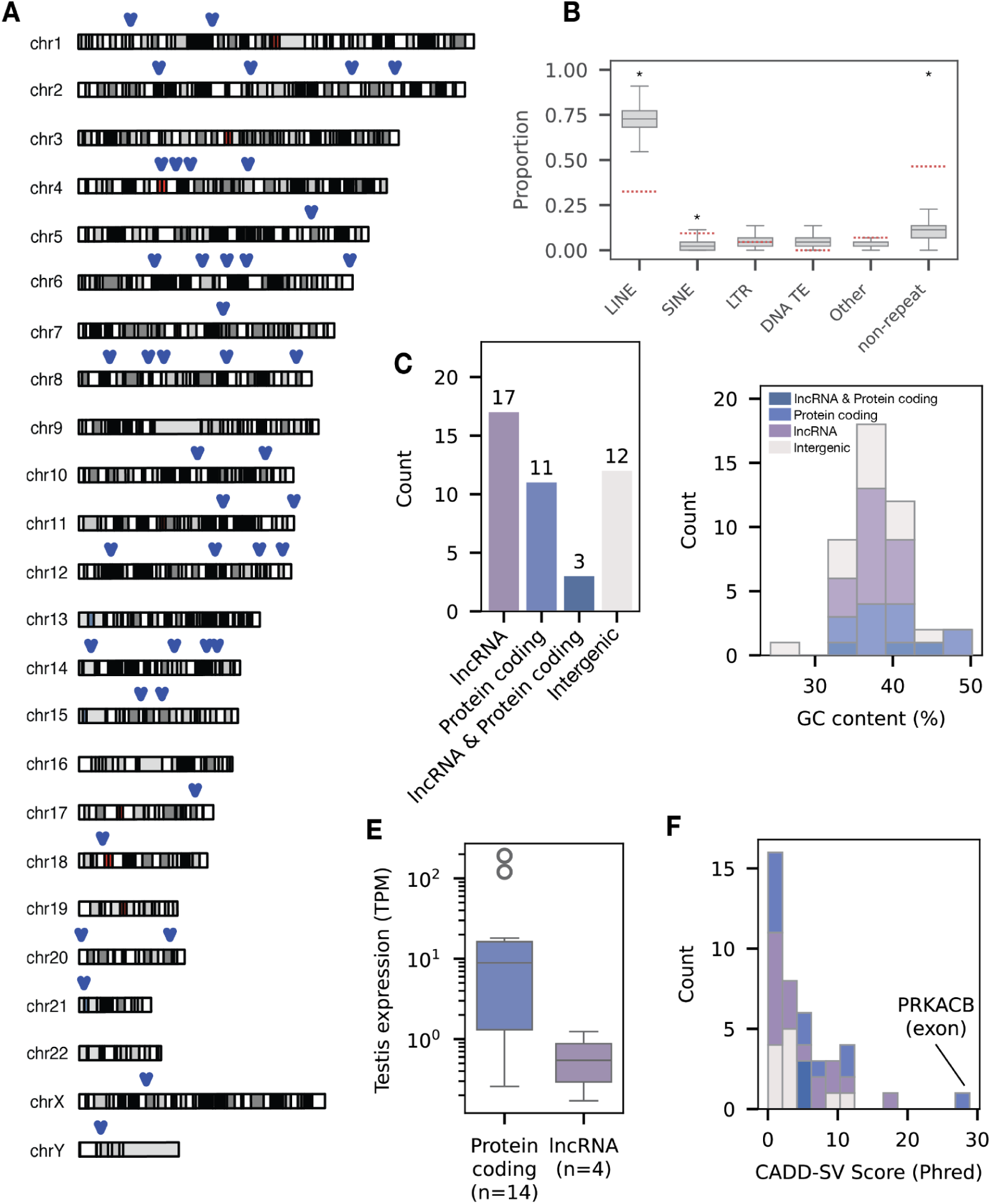
Sequence preferences of *de novo* Alus. **A)** Chromosome ideogram of Alu insertion sites transferred to CHM13 coordinates. **B)** Boxes show the expected distribution of the proportion of *de novo* Alu insertion sites across repetitive sequence annotations from RepeatMasker as well as non-repetitive sequence. Expected distributions were randomly resampled using degenerate L1 endonuclease motif coordinates. Red dotted lines show observed proportions of de novo Alu insertions across repeat features and non-repeats (* denotes p <0.05, FDR corrected permutation test). **C)** Distribution of GRCh38 insertion sites covered by GENCODE annotations. **D)** GC content in the 20 kb window around insertion sites stratified by intersected features. The average GC content is 38% across all insertions, compared to 41% observed across the total human genome. **E)** Gene expression from GTEx bulk RNA-seq analysis of testis tissue. All intersected genes are expressed in the testis. **G)** CADD-SV Phred-scaled scores for all GRCh38 insertion sites.

Interestingly, we find a significant enrichment of *de novo* Alu insertions in Alus (SINEs) but not in LINEs (Hoyt et al. 2022)(**Fig. 4B**; FDR-corrected two-sided permutation test P=0.014 for both). Instead, we observe a significant depletion of Alu insertions in LINEs compared to expectations, as well as a significant enrichment of Alus outside of repeats annotated by RepeatMasker (FDR-corrected two-sided permutation test P=0.003 for both; **Fig. 4B**). This discrepancy between our findings and some previous reports may reflect differences in null models for detecting enrichment. For example, (Levy et al. 2010) and colleagues did not incorporate expected L1 endonuclease integration site preferences into their null model when testing for enrichment of Alus in other repeats (Levy et al. 2010). We note that we do indeed find a >1.5-fold enrichment of *de novo* Alu insertions within LINEs when we consider a null wherein the expected probability of insertion is simply the proportion of the genome occupied by LINEs (Binomial test, P=0.047). These results highlight differences between the genomic distribution of *de novo* Alu insertions and the distributions of segregating and fixed Alus.

Both ancient and segregating Alus are enriched in GC-rich, gene-dense regions of the genome (Grover et al. 2004b; Jurka et al. 2002; Gu et al. 2000; Lander et al. 2001). To explore associations between *de novo* Alu insertions and genes, we projected Alu insertion coordinates from each personalized assembly to the GRCh38 reference genome, excluding two insertions without GRCh38-equivalent coordinates (remaining n=41). Then, we intersected *de novo* Alu insertions with protein coding and lncRNA gene features annotated by GENCODE. The majority (31/41) of Alus intersected at least one feature, with ∼34% (14/41) of insertions intersecting protein-coding genes and nearly half (20/41) intersecting at least one long non-coding RNA gene (lncRNA) (**Fig. 4C**). While the proportion of Alu insertions in genes is higher than expected, the difference is not statistically significant (75.6% observed vs 70% expected; Permutation test, P=0.18). Furthermore, the distributions of insertions in introns versus exons largely mirror their relative genomic composition. Nearly all (30/31) genic insertions occurred within introns, reflecting the minor fraction of gene content occupied by exon sequences (X^2^ test, P=0.882) (Francis and Wörheide 2017) with a single exonic insertion in the *PRKACB* gene. We also assessed the GC content around *de novo* Alu insertion sites and found that they are enriched in AT-rich regions (62%-AT compared to 59%-AT in the human genome on average; Permutation test, N=1000, P<0.001; **Fig. 4D**). This result likely reflects the AT-rich insertion site preference of the L1 endonuclease (Jurka 1997), and may explain the more random genomic distributions displayed by young Alus compared to more ancient Alu families (Cordaux et al. 2006b).

Biophysical simulations of chromatin accessibility suggest that retrotransposons preferentially integrate in open, sterically permissive chromatin regions (Prizak et al. 2025; Michieletto et al. 2019). Under this premise, we conducted an analysis of the GTEx bulk RNA-seq dataset, reasoning that genes expressed in the testis are reasonable proxies of chromatin accessibility. We retrieved GTEx data for 18/31 intersected genes, including all 14 protein-coding genes (**Fig. 4F**). All genes with Alu insertions are expressed in the testes (TPM > 0.1), suggesting that chromatin accessibility during spermatogenesis and in mature sperm may shape the genomic distribution of *de novo* insertions in the germline. Together our data supports a model wherein Alus exhibit an AT-rich preference but primarily insert into expressed genes likely due to their more permissive chromatin.

Finally, we used CADD-SV to conduct an exploratory functional analysis of each insertion (Kleinert and Kircher 2022). CADD-SV predicts and scores the effects of SVs using random forest models trained on variants from human and chimpanzee-derived SVs. Scores are Phred-transformed relative to variants of known phenotype from the gnomAD-SV dataset, ranging from 0 (unlikely pathogenic) to 48 (likely pathogenic): for example, a score of 10 corresponds to the 90th percentile of estimated pathogenicity. We scored all 41 variants, including intergenic variants, that were successfully projected to GRCh38 coordinates (**Fig. 4G**). Out of the six insertions with a score greater than 10, three occurred in protein-coding genes. As expected, the single exonic insertion in *PRKACB* received the highest score of any variant in the dataset. *PRKACB* is one of two catalytic subunits of *PKA*, a critical regulator of human morphogenesis. Germline and mosaic variants of *PRKACB* are associated with congenital malformations and neurodevelopmental disorders (Palencia-Campos et al. 2020). The second highest score corresponds to an intronic insertion in *Lnc-RUNX1T1-3*, a lncRNA with poorly understood functions.

## Direct estimation *de novo* Alu retrotransposition rates from gametes

Single molecule sequencing and *de novo* identification of Alus in gametes enables personalized estimates of the mutation rate. We calculated the per-sample, per-gamete *de novo* Alu retrotransposition rate as the number of observed Alu insertion events divided by sequencing coverage, as each read represents sampling of a single gamete. **(Fig. 5A)**. Combining our estimates across samples, we estimate an average rate of 4.52 Alu insertions per 100 gametes for all surveyed individuals (95% CI: 3.32-5.94 Alu insertions per 100 gametes), i.e. a mutation rate of 0.0452. This estimate falls within the distribution of mutation rates derived from pedigree and population studies of Alu activity which range from 0.018-0.07, highlighting congruence with population-based estimates (**Fig. 5B**) (Feusier et al. 2019; Hancks and Kazazian 2012; Hormozdiari et al. 2011; Werling et al. 2018; Cordaux et al. 2006a). The confidence intervals of our *de novo* mutation rates intersect that of previous estimated mutation rates. However, we note that our estimates only take into account the male germline while pedigree-and population-based approaches consider the combined male and female mutation rates.

**Figure 5:**
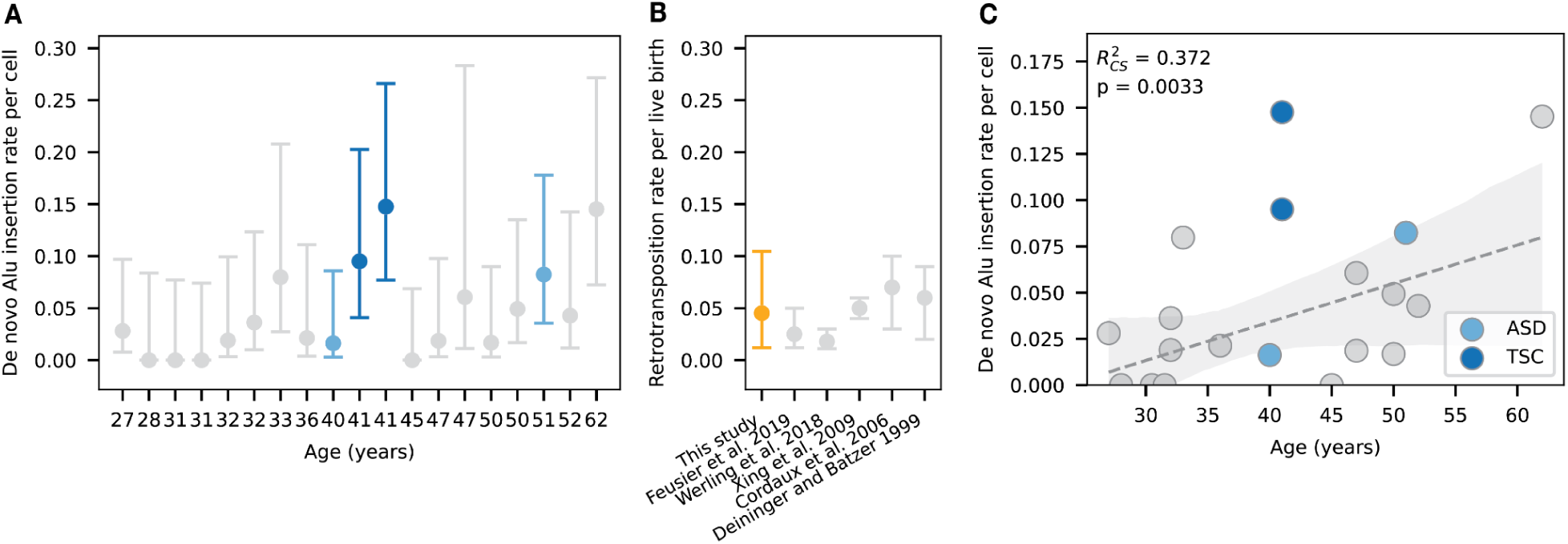
Individual and population-level Alu retrotransposition estimates. **A)** Per-individual de novo Alu retrotransposition rates by age. **B)** Aggregate *de novo* Alu retrotransposition rate compared to population genetic estimates. **C)** Scatter plot showing the association between *de novo* Alu insertion rate and donor age across human sperm samples. A generalized Poisson regression model between donor age and insertion rate is shown with 95% confidence intervals. Donors with offspring affected by ASD and TSC were excluded when fitting this model. The Cox and Snell pseudo R-squared (*R^2^*_cs_) and p-value for this model are displayed in the top left corner. Individuals with offspring affected by ASD (autism spectrum disorder) and TSC (tuberous sclerosis complex) are highlighted. We note that the two points for 31 year old individuals with *de novo* Alu insertion rates of zero are jittered.

Next, we examined covariation between age and Alu insertion rates. We observe a >9-fold range in rates estimated among individuals in our dataset, ranging from 0 to 0.148 *de novo* Alus per gamete. Notably, we identify a statistically significant increase in Alu retrotransposition with age in healthy samples, estimated at an increase of 4.78% Alu insertions per gamete per year of paternal age (p = 0.0033, *R^2^_cs_*= 0.372, linear regression) **(Fig. 5C)**. We also examined Alu retroposition rates in samples from donors with offspring affected by ASD (n=2, “ASD samples”) and TSC (n=2, “TSC samples”). We did not observe a significant difference in the mutation rates of ASD samples. However, we identify an elevated mutation rate in one of our TSC samples (p = 0.0036, two-tailed z-test). TSC is a neurodevelopmental disease predominantly caused by *de novo* mutations in the *TSC1* and *TSC2* genes. In several cases, TSC has also been shown to be caused by paternal gonadal mosaicism (Verhoef et al. 1999; Uysal and Şahin 2020). However, we found no evidence of Alu insertions nor mosaicism in *TSC1* or *TSC2* in either TSC sample.

## Discussion

Here we demonstrate how long read gamete sequencing can be used to characterize *de novo* mobile element mutation processes in the human germline at unprecedented scale and resolution. Previous work has employed similar methodology using short reads (Freeman et al. 2011). By optimizing variant calling methods for ultra-low frequency variant detection, we directly isolate *de novo* retrotransposition events without the use of family-based inference.

We demonstrate the feasibility of constructing high-quality genome assemblies from haploid sperm samples. Self-alignment to personalized genome assemblies addresses a persistent technical challenge in detecting low-frequency genetic variation: distinguishing true variation from artifacts arising during sequencing and alignment. Simulations demonstrate that self-alignment substantially improves sensitivity compared to standard reference alignment. This increase in sensitivity underlies the feasibility of single-read and low allele fraction SV discovery where read depth approaches are not applicable.

We characterize 43 individual *de novo* Alu insertions across 19 individuals, identifying detailed molecular features of each insertion and its genomic sequence context. The subfamily composition of our *de novo* insertions is highly consistent with patterns observed in segregating Alu elements from the 1000 Genomes Project, with AluYa5 and AluYb8 subfamilies comprising the majority of insertions. This concordance validates both direct observation and population genetic approaches while demonstrating that current retrotransposition activity reflects the same processes that generated segregating variation in recent evolutionary history. Furthermore, we capture the full span of each element’s poly(A) tail directly in long reads. These tails are significantly longer than those observed in ancient, immobile elements, supporting models of progressive tail degradation following insertion. In contrast to these observations however, reporter assays in HeLa cells exhibit substantially longer poly(A) tails than observed in the germline (Wagstaff et al. 2012), potentially suggest differences between somatic and germline TE insertion mechanisms which we expect will be illuminated by future experiments to directly profiling somatic TE insertions using long reads.

We directly estimate *de novo* mobile element insertion rates for each individual, providing insights into inter-individual variability in germline mutation rates. We find a significant association between MEI rate and donor age in healthy samples, reflected in the >9-fold difference across the donors in our study. Several mechanisms may account for increased TE mutation rates in the male germline. If TE insertion occurs stochastically in the germline at a constant rate then an age-associated increase would be expected. However, age-associated dysregulation of other molecular processes may also contribute. Age-associated changes in chromatin might result in a DNA landscape that is more permissible to TE insertion. For example, several DNA methyltransferase enzymes are essential for the suppression of young TEs in the male germline (Barau et al. 2016). Age-associated changes in the expression of these epigenetic regulators could impact TE insertion rates. Similarly, age-associated dysregulation of the PIWI-piRNA pathway could impact TE activity. L1 source elements, which encode for the ORF2p endonuclease, are typically repressed via hypermethylation and post-transcriptional degradation by the PIWI-piRNA pathway (Gainetdinov et al. 2017; Vagin et al. 2006). Studies in both mouse models and clinical cases of infertility suggest that dysregulation of piRNA biogenesis may result in impaired spermatogenesis and de-repression of L1 elements in early meiosis (Wei et al. 2023; Stallmeyer et al. 2024). Another hypothesis is that TE insertion is coupled with other processes of DNA damage and repair. Increasing evidence suggests that *de novo* single nucleotide mutations in both the female and male germline are the result of the continuous impact of DNA damage and repair processes (de Manuel et al. 2022). Such damage or repair processes could potentially provide opportunities for TE insertions, particularly if they are associated with double stranded breaks (DSBs). An increase in the number of DSBs with age has been observed in sperm (Laurentino et al. 2020). Anecdotally, we observe a significantly elevated mutation rate in a sample provided by a donor with a child affected by tuberous sclerosis complex, suggesting potential associations between individual mutation burden and disease risk that merit further investigation.

Finally, we repeatedly identify close agreement between our direct measurements of retrotransposition rates and estimates derived from population genetic, and pedigree-based studies. This concordance suggests that the events we observe in gametes are representative of those that successfully transmit through the germline. This agreement also supports the use of segregating element analysis for understanding historical retrotransposon activity and evolution of retrotransposon lineages.

Our study has several notable limitations. In particular, we rely on statistical estimation of unique reads per cell, precluding direct identification of cell-specific mutations and reconstruction of clonal mutations. Single-cell approaches, while technically more challenging, could provide higher resolution insights into the timing and cellular context of mutagenic events. Additionally, due to the destructive nature of the sampling and sequencing process, we lack the power to deeply follow up on the mechanisms underlying age-associated increases in the rate of TE insertion. For the same reasons, we further lack the ability to validate any individual *de novo* insertion. Another limitation of our approach is that we require the presence of TSDs for accurate filtering of *de novo* MEI candidates. TSDs are a hallmark of most known TEs, including all transpositionally competent TEs in humans (Cordaux and Batzer 2009; Wells and Feschotte 2020). However, there are some known TEs that fail to generate TSDs upon insertion such as rolling circle transposons and cryptons (Wells and Feschotte 2020). This may affect pipeline sensitivity in some species divergent from humans. Nevertheless, we expect this pipeline will have extensive utility to explore the dynamics of *de novo* TE insertions across diverse species.

Overall, we demonstrate that direct sequencing of gametes from even a comparatively small population of individuals has the power to accurately quantify the mutation rate of mobile elements. We identify a significant age-associate increase in the mutation rate of *de novo* structural variants. Our results demonstrate the feasibility of using highly accurate long-read sequencing to directly observe germline mutation at the individual level. Future studies utilizing direct gamete sequencing will improve our understanding of structural variants, individual variation in germline mutation rates, and genetic disease risk.

## Methods

### Study design and sample collection

We obtained semen samples from donors from ages 27 to 62, with a median age of 41. Four samples from fathers of children with autism spectrum disorder (ASD, n=2) and tuberous sclerosis complex (TSD, n=2) were shared by the Gleeson lab. All samples were stored at-80°C until processing. Discarded medical waste samples were obtained from Discovery Life Sciences in accordance with IRB protocol DLS-BB050. All samples were collected as part of standard-of care testing ordered by the patient’s physician and excess material released for research.

### DNA extraction

Semen samples were thawed on ice and subjected to a gradient separation using 90% PureCeption Isotonic Solution (CooperSurgical Cat. #ART-2100, Cat. #84051010) to remove seminal fluid and contaminating somatic cells. After separation, the cell pellet was washed with 2 mL of PBS and resuspended in 100 uL of PBS in a 2 mL centrifuge tube. DNA extraction was performed using a modified version of the Monarch® HMW DNA Extraction Kit for Cells & Blood (New England Biolabs Cat. #T3050L) extraction protocol. Cell lysis solution (100 uL of Nuclei Prep Buffer, 100 uL of Lysis Buffer, 15 uL of 20 mg/mL Proteinase K, 15 µL of 1M DTT (Goldbio Cat. #DTT)) was prepared and added to each sample, followed by thermal mixing with an Eppendorf ThermoMixer® set to 56°C/300 RPM for 2 hours. RNA digestion was performed by adding 5 uL of 20 mg/mL RNase A and incubating under the same conditions for 20 minutes. The remainder of the DNA binding and wash steps were conducted according to the protocol description. DNA was eluted from glass beads overnight at room temperature using 75 uL of Elution Buffer, followed by centrifugal separation into Eppendorf DNA LoBind® Tubes (Eppendorf Cat. #0030108523). Sample DNA was gently mixed by wide bore pipetting and then allowed to rest for 24-48 hours at room temperature before quantification using the Promega® Quantus™ Fluorometer and purity ratio (A260/280, A260/230) assessment with a ThermoFisher NanoDrop Spectrophotometer. Samples were submitted to the UC Berkeley Functional Genomics Laboratory for fragment analysis using an Agilent Femto Pulse system. Samples containing at least 10 µg of DNA with a median fragment length exceeding 50 kB were deemed sufficient for sequencing. DNA was stored at 4°C and shipped on dry ice to the sequencing laboratory.

### Long-read sequencing

Highly accurate long-read whole-genome DNA sequencing was performed at the HudsonAlpha Genome Sequencing Center using the Pacific Biosciences (PacBio) Revio sequencer. We sequenced each sample using two SMRTcells to a target depth of 50X haploid genome coverage with an 18 kb read length, yielding unaligned BAM files for each SMRTcell. Samples with insufficient coverage were resequenced to a maximum of four total SMRTcells. We converted each uBAM to FASTQ format using samtools (v.1.21) (Li et al. 2009), preserving all tags in the read headers. We used HiFiAdapterFilt (Sim et al. 2022) to confirm the absence of errant sequencing adapters present in data from previous generation sequencing machines.

### Estimating reads per unique cell

We estimated the maximum number of reads per unique cell using 50X coverage and a normally distributed read size (mean = 18 kb, standard deviation = 2.5 kb), representative of the average values for each sample. Each sample of sperm cells is subsampled from the population of gametes produced by the individual. We assume that this population is large, each meiotic cycle is perfect (produces 4 daughter cells), and all 4 daughter cells are represented in the population. The average sperm cell count and volume in each semen sample is 2 mL of 100 million cells/mL per specimen, for a total of 200 million cells representing approximately 50 million meioses.

From this specimen, we sample approximately 2.5 million cells for DNA extraction. Each sample was sequenced with 2 SMRTcells, each requiring 293 ng of DNA input.

To calculate maximum possible per-cell coverage, we assume a perfect yield of 18 kb fragments for each cell,

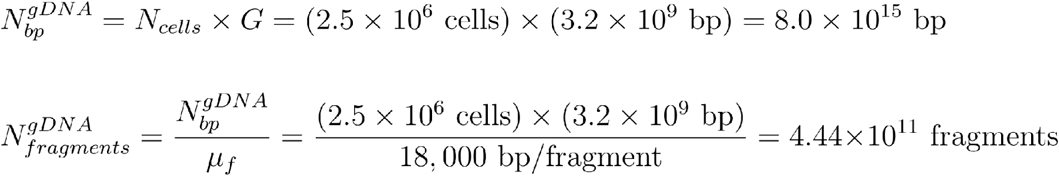

We then use the mass of DNA loaded for sequencing to calculate the number of prepared fragments for sequencing.

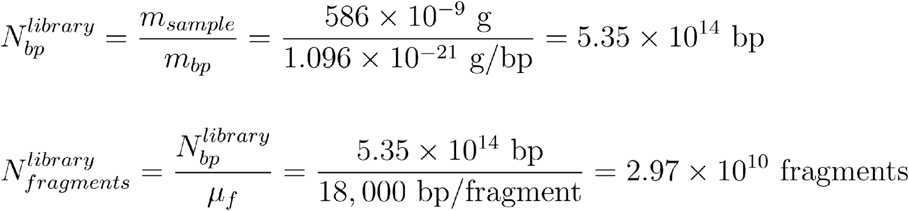

Knowing that our final coverage is 50X, we estimate *N_reads_^sequenced^* sequenced from *N_bp_^sequenced^*:

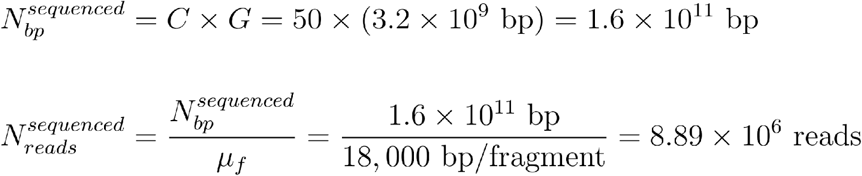

Finally, we calculate the maximum number of fragments sequenced per cell:

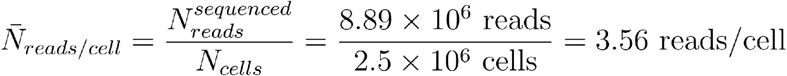

### Modeling unique meioses represented per sample

As described in the previous sections, we extracted DNA from approximately 2.5 million cells per specimen, such that the sampling probability per cell is:

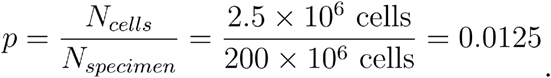

For each meiotic event, the number of its cells appearing in the sample follows a hypergeometric distribution. However, given the large population of cells in the specimen, we can accurately approximate this with a binomial distribution. For a single meiotic event with 4 cells, let *X* be the number of cells sampled, where:

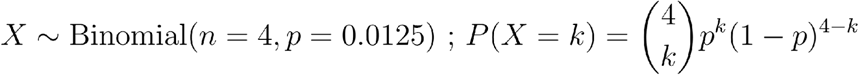

We calculate the probability of sampling a single daughter cell per meiosis (“singleton”) as:

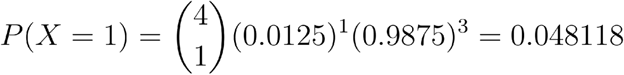

Thus, for 4.81% of all 50 million meioses, we obtain singletons totaling 96.2% of the cells in the sample.

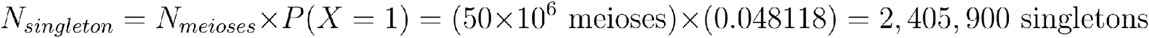

## Analysis

### Genome assembly

We generated haplotype-resolved assemblies using hifiasm (v0.25.0-r726) (Cheng et al. 2021) with default parameters. We did this for each sample presented in this study as well as an additional diploid sample for assembly comparisons (HG002). The primary raw assembly graphs (.gfa) for each haplotype were converted to FASTA formatted contigs using gfatools (v.0.5) (Li) and assessed using QUAST (v.5.2.0) (Gurevich et al. 2013). The average N50 of each assembled haplotype was 71.12 Mb, comparable to the N50 of the HG002 assembly haplotypes. Contigs were scaffolded to the T2T-CHM13-v1.0 reference genome using RagTag (v.2.1.0) (Alonge et al. 2022) and merged into a final FASTA file containing both haplotypes.

### Sequence alignment

We used minimap2 (v.2.26) (Li 2018) with parameters-ax map-hifi-Y-y-L --eqx --cs-I8g --MD to align HiFi reads for each sample against both the sample’s own assembly (“self-assembly”) and the T2T-CHM13-v1.0 reference genome for comparative purposes. We sorted, merged, and indexed resulting bam files for each sample using samtools (v.1.21) (Li et al. 2009). Then, we used deeptools (v.3.5.2) to query read depth, assess read alignment qualities, and calculate genome-wide coverage for each sample prior to variant calling (Ramírez et al. 2014).

### Genome annotation

We generated *de novo* tandem and low-complexity repeat annotations for each self assembly using ULTRA (v1.2.1) (Olson and Wheeler 2024) and longdust (v1.3-r87) (Li and Li 2026) with default parameters. These repeat annotations are important for filtering and clustering during downstream mosaic SV calling steps (Smolka et al. 2024; Qin et al. 2025; Qin and Li 2025).

### Variant calling

We used *sniffles2* (v.2.5.2) (Smolka et al. 2024) in mosaic mode with flags --output-rnames --mapq 0 --minsupport 0 --mosaic-af-min 0 --mosaic-af-max 0.1--mosaic-qc-strand False --dev-no-qc to raise a “first pass” callset of SV candidates. We optimized for recall of *de novo* events by lowering the minimum read support to a single read while also limiting the allele frequency of calls to no more than 0.10. We used the self-assembly as the reference for each sample and provided assembly-specific repeat annotations for repeat aware clustering (Smolka et al. 2024). We also applied the --dev-no-qc option to emit all variants regardless of the FILTER flag. We obtained supporting read names using the --output-rnames option. To evaluate sniffles2’s performance against other recently developed tools, we also called variants with miniSV (vr29) (Qin et al. 2025). Similar to our approach with sniffles2, miniSV employs self-assemblies to call low-frequency mosaic and *de novo* SVs from bulk pacbio data. MiniSV requires alignments to both a reference genome and self-assembly for reliable mosaic SV calling. We generated these alignments using minimap2 with miniSV-developer recommmended flags-cxmap-hifi-s50 --ds. To call low-frequency mosaic SVs, we ran minisv e with flag-0b and provided both alignment files as well as the low-complexity repeat annotations generated for each self-assembly. We merged resulting SV calls using minisv merge with flags-c1-s0-e1 and filtered for insertions occurring in less than 2 reads. Then, we generated a vcf from these calls using minisv genvcf.

### Pipeline for identifying de novo mobile element insertion

We developed a pipeline to identify *de novo* mobile element insertions from low-frequency variant calls in VCFs. The pipeline requires a VCF as input. First, we employed bcftools filter (v1.19) (Danecek et al. 2021)to filter a VCF for insertions >100 bp and <16kb in length with a minimum supporting coverage of 9 and a maximum allele frequency of 0.1. Coverage = 9 is the minimum coverage for which we can confidently distinguish low frequency variants from random expectations (two-tailed binomial test, p = 0.0390). We also required that insertions were spanned entirely by at least one Pacbio read. Then, we applied a sequence of stringent downstream filters to accurately distinguish true *de novo* mobile element insertions (MEIs) from false positives and noise. True nonLTR retrotransposon insertions (e.g. L1, Alu) should exhibit homology to their consensus sequences across the entire insertion sequence, should be mostly devoid of low-complexity sequences other than their poly(A) tail, and should exhibit identical target site duplications (TSDs) at their flanks. We retrieved a curated set of human and ancestral mobile elements from DFAM. Then, we used GraffiTE’s (v1.1dev commit 19b8090) (Groza et al. 2024) repeat filtering and TSD modules to annotate insertions for TE homology and TSDs as well as to filter for high-confidence insertions. Briefly, GraffiTE annotates VCFs by scanning insertion or deletion alleles with RepeatMasker (from a TE library) and ULTRA (for low complexity regions); it further retains SVs for which repeat annotation spans >= 80% of the sequence before searching for TSDs. We provided our filtered VCF and curated the TE library to GraffiTE with parameter --genotype false. Then, we further filtered GraffiTE’s output VCF for insertions >250 bp, with n_hits=1, and that were not primarily Satellites per the developer’s benchmark (Groza et al. 2024). We also filtered for MEIs exhibiting flanking TSDs and <3% Kimura Divergence from their consensus, as expected for recently inserted mobile elements (Konkel et al. 2015; Neafsey et al. 2004; Bennett et al. 2008). We additionally filtered for MEIs with poly(A) tails, which are required for L1 and Alu transposition (Dewannieux and Heidmann 2005b; Doucet et al. 2015)e reasoned that the probability of observing identical *de novo* MEIs multiple times with the same TSDs on the same chromosomes is low, and that any MEIs meeting these criteria are likely false positives due to contamination or mapping artifacts in highly homozygous regions. To identify these potential false positives, we blasted the sequence of each MEI with its flanking TSDs to its corresponding self-assembly and CHM13 using blastn (v2.17.0) (Morgulis et al. 2008)

### Pipeline evaluation on simulated data

We developed a simulation framework to benchmark our approach for *de novo* MEI identification as well as the effects of variant callers and choice of reference on its performance.

This simulator, which we named *nova*, introduces predefined variants to individual reads from a reference dataset. For each human sperm sample, we used *nova* to simulate 500 random insertions of length 250 bp, 500 random insertions of length 1 kb, 600 Alu insertions, and 200 L1 insertions in single reads sampled from bulk Pacbio data. We used random insertions in single reads as negative controls and synthetic MEIs as positive controls. Synthetic Alu and L1 insertions were generated using the consensus sequences for known families in humans as well as partial elements in the case of L1 (exact config file: https://github.com/sudmantlab/nova/blob/main/examples/Li_et_al_params.yml). We also simulated TSDs of length 6-20 bp at the expected positions for each synthetic TE insertion.

Then, we used these synthetic reads to benchmark our pipeline. First, we used sniffles2 mosaic for variant calling and compared pipeline performance when mapping to the T2T-CHM13-v1.0 reference genome versus mapping to self-assemblies, finding that mapping to sample-specific self-assemblies dramatically improved sensitivity as well as accuracy (Supplementary Figure 1). We also compared pipeline performance between sniffles2 mosaic and another recently developed mosaic structural variant caller miniSV. We found that sniffles2 was both more sensitive and more accurate than miniSV. Thus, we implemented sniffles2 in our finalized workflow. Notably, our pipeline produced zero false positives in all benchmarks, irrespective of the upstream variant caller or reference, suggesting robust automated curation of prospective *de novo* MEIs.

### Identifying and filtering prospective de novo mobile element insertions in human sperm samples

We applied our de novo MEI pipeline to sniffles2 mosaic self-assembly callsets for each human sperm sample. This produced a total of 45 candidate *de novo* Alu insertions and 1 candidate *de novo* L1 insertions. We also manually inspected all candidate MEIs for expected sequence features such as poly(A) tails and TSDs. We also inspected read coverage across each prospective *de novo* insertion and its flanking regions. Changes in relative coverage across structural variants could suggest potential mapping artifacts (Smolka et al. 2024). We visualized each insertion in IGV (Robinson et al. 2011) (see https://github.com/sudmantlab/denovo-TE-insertion-discovery/tree/main/paper/igv for IGV html files). Manual inspection of *de novo* MEI candidates revealed one potential false positive Alu insertion with low mapping quality within a segmental duplication. All remaining 44 MEIs passed manual inspection. Next, we lifted over each VCF to the T2T-CHM13-v1.0 reference genome using paftools liftover (Li 2018) and merged callsets across samples using SURVIVOR with parameters 1000 0 1 0 0 80 (Jeffares et al. 2017). Then, we sought to further evaluate potential false positives due to contamination. We downloaded an SV callset generated from 1,019 long-read human samples as part of the 1000 Genomes Project (Schloissnig et al. 2025). Then, we flagged any de novo MEIs whose insertion site was within 50 bp and whose length differed by ≤50 bp from any MEI in the 1000 Genomes callset. We identified only one candidate *de novo* Alu insertion meeting these criteria, suggesting that our aforementioned filters are robust for removing false positives from contamination. After removing this potential false positive, our final set of high-confidence *de novo* MEIs included 43 Alus and a single L1. Due to the low number of L1s, we focused our downstream analyses on *de novo* Alu insertions.

### Analysis of ORF2p motif, target site, and extended context

We performed the proximal insertion context analysis by extracting the 4 bases surrounding the insertion breakpoints. For sense-oriented Alu insertions, we extracted the 4 base pairs upstream of the “left” TSD start coordinate and concatenated them to the 4 base pairs following the poly(A) tail end coordinate. For antisense-oriented Alu insertions, we extracted the 4 base pairs upstream of the poly(A) tail end coordinate and concatenated them to the 4 base pairs following the “right” TSD end coordinate. We used Logomaker through python to generate insertion site sequence logo plots (Tareen and Kinney 2020).

### Insertion site context and functional implication analysis

We used karyoploteR to plot the ideogram in Figure 5 (Gel and Serra 2017) To account for the effects of segregating genetic variation, we analyzed the GC content and genomic repeat features at *de novo* Alu insertion sites in the context of their respective self assembly. To assess GC content, we used bedtools slop (v2.29.1) (Quinlan and Hall 2010) to create a 20 kb window around each Alu insertion. Then, we calculated GC content using bedtools nuc. We performed a permutation test for enrichment of Alu insertions in GC or AT-rich genomic regions. Specifically, we compared the observed mean GC-content around Alu insertion sites to equally sized windows randomly resampled from the CHM13 reference across 1000 permutations. To intersect Alu insertions with genomic repeats, we first generated interspersed repeat annotations for each self-assembly using RepeatMasker (v4.2.1) (https://www.repeatmasker.org) with flags-excln-e ncbi-s-no_is-u-noisy-html-xm-a-xsmall-source and the aforementioned curated human TE library as input. We converted RepeatMasker output files to bed format using the RepeatMasker utility tool RM2Bed.py and intersected these annotations with *de novo* Alu insertion sites using bedtools intersect. Then, we performed a two-sided permutation test comparing the proportion of Alu insertions in different repeats to expectations from random resampling of degenerate L1 motif coordinates using the approach and models described in (Flasch et al. 2019) (N=1000). We performed a false discovery rate correction on resulting P-values. To explore the possible pathogenic implications of Alu insertions, we lifted over our Alu callset to HG38 using UCSC’s online liftover tool (Casper et al. 2026). Two Alu insertions could not be confidently placed and were filtered from downstream analyses. We used bedtools intersect with annotation files from GENCODE (v50) to identify genic features present at insertion loci and assess the intron/exon boundaries for each feature. Then, we used the aforementioned approach in which we random resampling of degenerated L1 motif coordinates to test for enrichment of Alu insertions in genes. We used Ensembl gene IDs to query GTEx (v10) median gene expression (TPM) values for each intersected gene. We used CADD-SV’s online submission system (https://cadd-sv.bihealth.org/) to run CADD-SV (v2.0) (Kleinert and Kircher 2022) on all *de novo* Alu insertions.

### Testing for associations between age and mutation rate

We used a Gaussian generalized linear model (GLM) through python’s statsmodels module to test for a correlation between donor age and mutation rate across human sperm samples (Seabold and Perktold 2010). We only used samples with healthy offspring in this analysis to control for the possible effects of disease on sperm mutation rate. We performed a two-tailed Z-test to test for significant differences in mutation rates between samples from individuals with children exhibiting TSC or ASD and other samples. Only TSC7237 differed significantly from the rest of our data.

## Declaration of Interests

The authors declare no competing interests.

## Acknowledgements

This work was supported by NIH National Institute of General Medicine award R35GM142916 and R01AG087974 to PHS. Sample collection was supported in part under NIH/NICHD R00HD111686 to XY. LG is supported in part by T32AG000266-27 in collaboration with The Buck Institute for Research on Aging and a University of California Chancellor’s Postdoctoral Fellowship. This manuscript is the result of funding in whole or in part by the National Institutes of Health (NIH). It is subject to the NIH Public Access Policy. Through acceptance of this federal funding, NIH has been given a right to make this manuscript publicly available in PubMed Central upon the Official Date of Publication, as defined by NIH. The content is solely the responsibility of the authors and does not necessarily represent the official views of the National Institutes of Health.

## Author Contributions

PHS conceived of the study. SL, LG, CC, and CG performed analysis. PHS, LG,SL, and CC wrote the manuscript. PHS, LG, SL, and CC edited the manuscript. KA, JG, AQ, and XY provided samples.

## Data availability

Supplementary Table 2 contains the raw read data supporting each insertion. Complete sequencing data for samples are not permitted to be deposited in searchable databases due to how they were consented; however data can be shared directly upon request to. The full analysis pipeline is available at https://github.com/sudmantlab/denovo-TE-insertion-discovery. *nova* is available at https://github.com/sudmantlab/nova.

## Supplementary Tables

**Supplementary Table 1:** Sequencing coverage of sperm samples considered in this study.

**Supplementary Table 2:** This table contains detailed information for each Alu insertion identified in this study. Columns include: sample (sample identifier), variant_id (unique variant identifier), variant_chrom (chromosome), variant_position (genomic coordinate), ins_name (Alu subfamily classification), original_strand (genomic strand orientation), ins_start_in_read and ins_end_in_read (insertion boundaries within the supporting read), ins_seq_length (insertion length in bp), ins_seq (inserted nucleotide sequence), tsd_seq (target site duplication sequence), tsd_length (target site duplication length in bp), left_tsd_start_relative_to_insertion, left_tsd_end_relative_to_insertion, right_tsd_start_relative_to_insertion, and right_tsd_end_relative_to_insertion (TSD positions relative to insertion), poly_a_tail_seq (poly(A) tail sequence), poly_a_tail_length (poly(A) tail length in bp), poly_a_tail_impurity (proportion of non-adenine bases in poly(A) tail), poly_a_tail_start_relative_to_insertion and poly_a_tail_end_relative_to_insertion (poly(A) tail positions relative to insertion), read_name (supporting read identifier), and read_seq (full supporting read sequence).

## Supplementary Figures

**Supplementary Figure 1:**
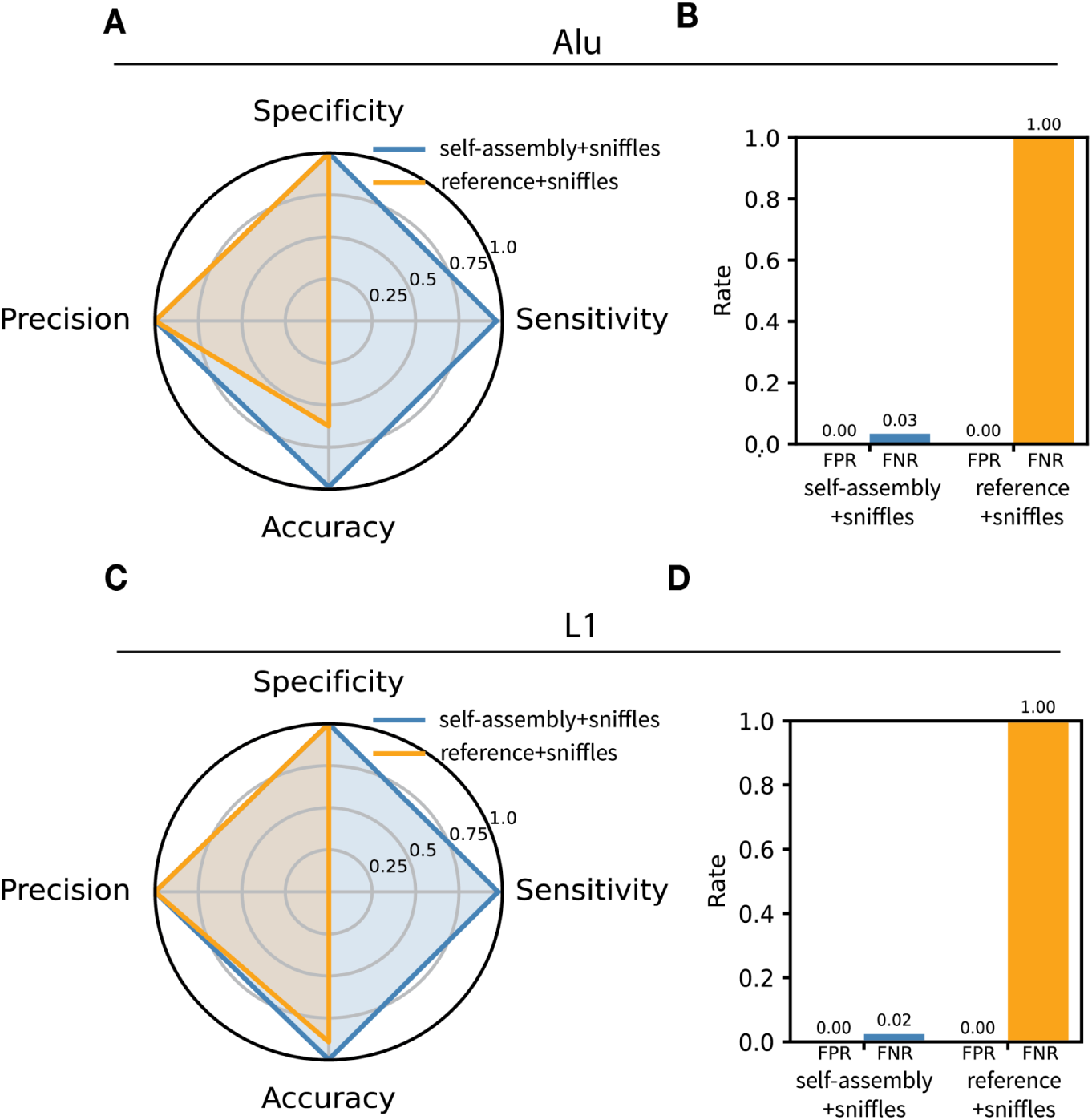
Comparison of *de novo* MEI detection performance between sniffles2 mosaic when mapping to self-assemblies versus the CHM13 reference. A) Radar plot comparing precision, specificity, accuracy, and sensitivity across self-assembly and reference alignment approaches for simulated *de novo* Alu insertions. B) False positive rates (FPR) and false negative rates (FNR) for simulated *de novo* Alu insertions. C) Radar plots comparing across self-assembly and reference alignment approaches for simulated de novo L1 insertions. D) FPR and FNR for simulated de novo L1 insertions.

**Supplementary Figure 2:**
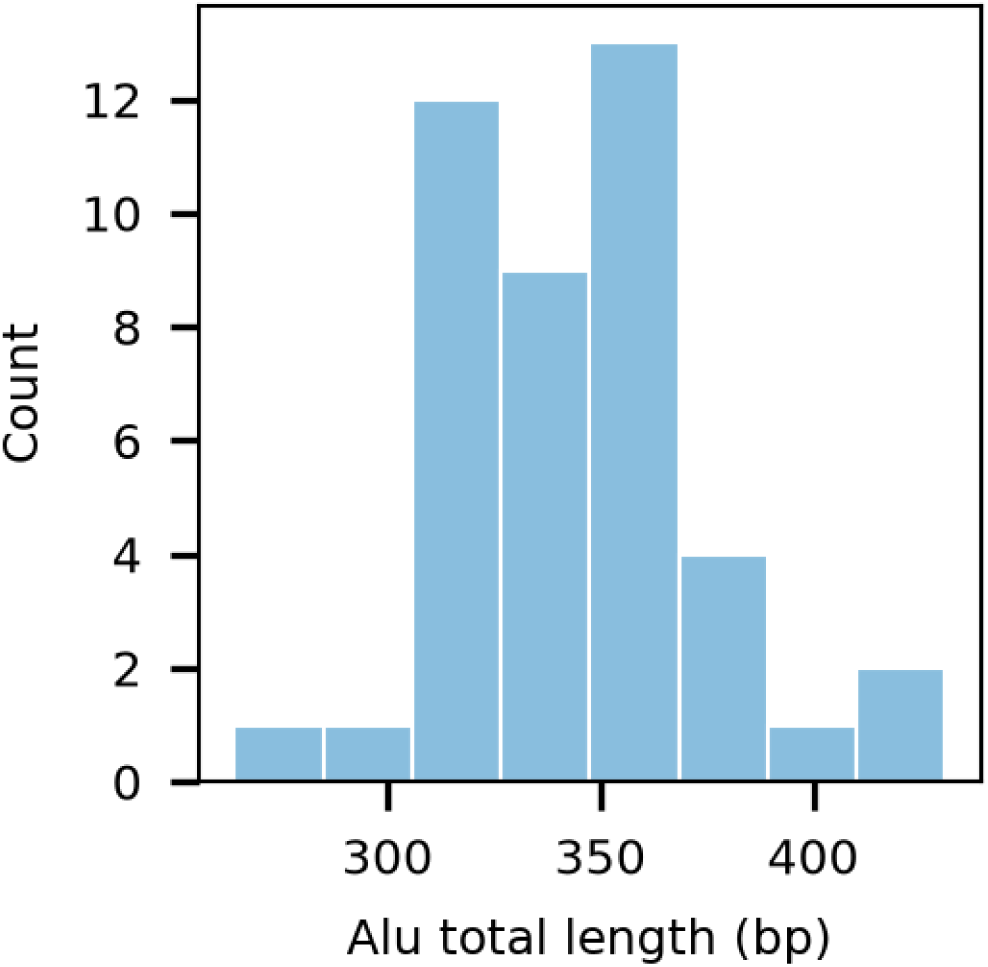
De novo Alu insertion length distribution (Alu body + poly(A) tail)

**Supplementary Figure 3:**
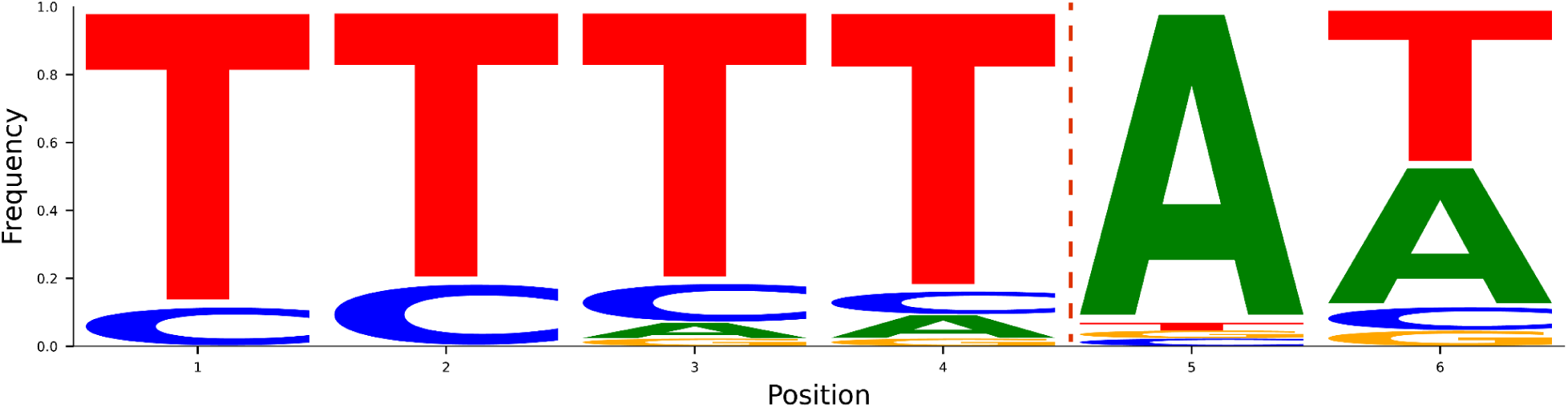
Sequence logo plot showing the integration site motif 5’-TTTT/AA-3’. The Alu integration site is shown with a dotted line.

**Supplementary Figure 4:**
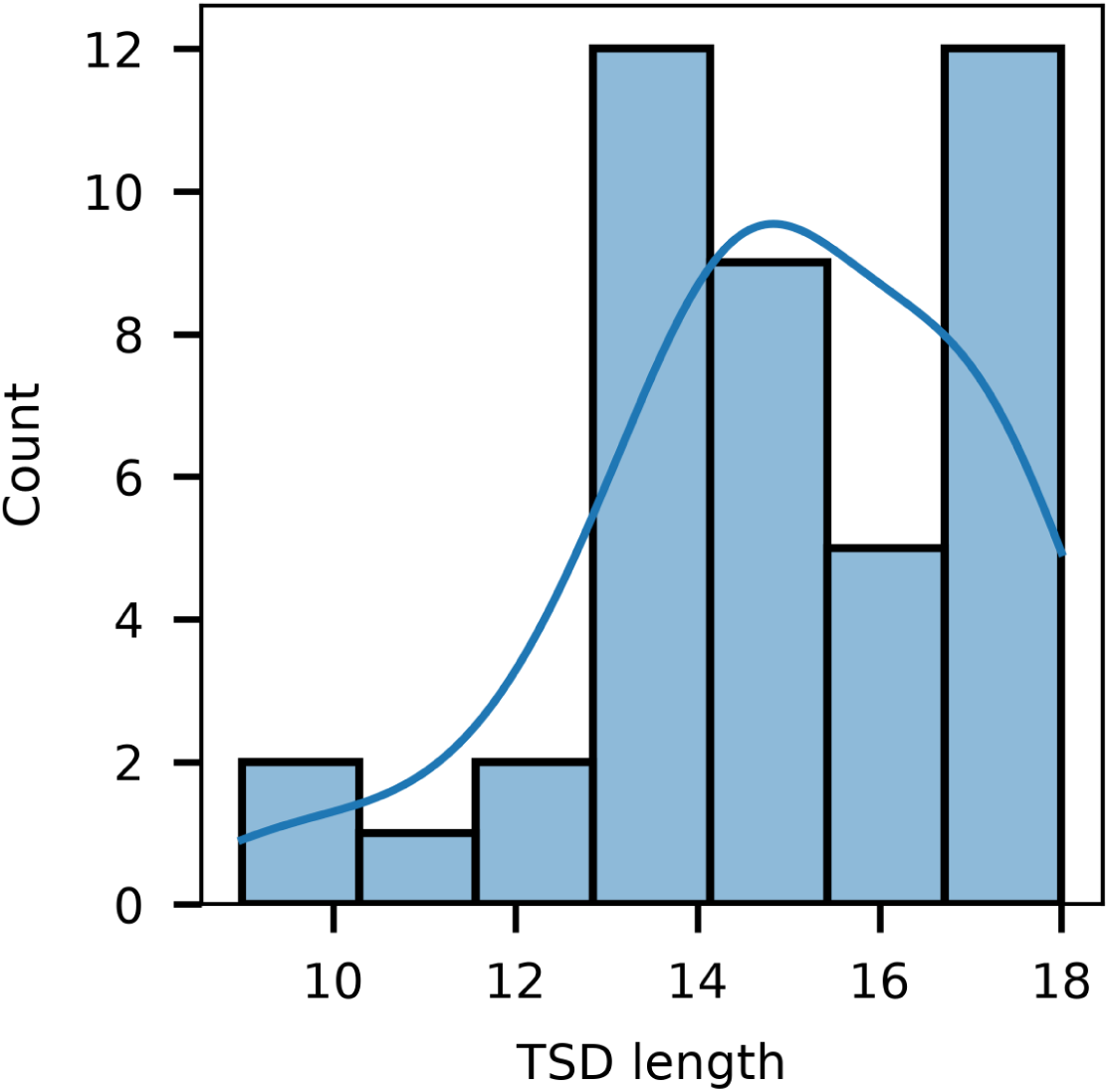
Distribution of TSD lengths for *de novo* Alu insertions. The blue line shows a kernel density estimate for the distribution.

## Notes

### Competing Interest Statement

The authors have declared no competing interest.

### Summary of Updates

This version of the manuscript has been revised to address reviewer comments and includes a fully updated pipeline which has improved precision and recall.

